# Genetic load in incomplete lupus erythematosus

**DOI:** 10.1101/2022.02.14.480411

**Authors:** Matt Slief, Joseph Kheir, Miles Smith, Colin Mowery, Susan Macwana, Wade DeJager, Catriona A. Wagner, Teresa Aberle, Judith A. James, Joel M. Guthridge

**Affiliations:** Arthritis and Clinical Immunology, Oklahoma Medical Research Foundation, Oklahoma City, OK, 73104; College of Medicine, University of Oklahoma Health Sciences Center, Oklahoma City, OK, 73104; Department of Medicine, Pathology, University of Oklahoma Health Sciences Center, Oklahoma City, OK

**Keywords:** systemic lupus erythematosus, incomplete lupus erythematosus, genetic load

## Abstract

Incomplete lupus erythematosus (ILE) patients have lupus features but insufficient criteria for systemic lupus erythematosus (SLE) classification. Some ILE patients transition to SLE, but most avoid major organ involvement. This study tested whether the milder disease course in ILE is influenced by reduced SLE-risk allele genetic load. We calculated the genetic load based on 99 SLE-associated risk alleles in European American SLE patients (≥4 ACR-1997 criteria, n=171), ILE patients (3 ACR-1997 criteria; n=174), a subset of ILE patients not meeting SLICC classification (ILE^SLICC^, n=119), and healthy controls (n=133). ILE and SLE patients had significantly greater SLE-risk allele genetic load compared to healthy controls, while ILE^SLICC^ patients had a trend toward an increased genetic load, although not statistically significant. However, the genetic load did not differ between ILE and SLE. In conclusion, ILE and SLE patients have comparable genetic loads of SLE risk loci, suggesting similar genetic predisposition between these conditions. Phenotypic differences between SLE and ILE may instead be influenced by ILE-specific variants and gene-environment interactions.

## INTRODUCTION

Systemic lupus erythematosus (SLE) is a complex chronic autoimmune disease with various systemic manifestations. SLE is typically diagnosed based on characteristic clinical and serological features defined by the American College of Rheumatology (ACR) or Systemic Lupus International Collaborating Clinics (SLICC)^1-3^. However, a subset of patients, referred to as incomplete lupus erythematosus (ILE), exhibit some clinical symptoms or serological evidence of SLE but do not fulfill classification criteria. Approximately 20% of patients with ILE transition to classified SLE within 5 years of onset, but most experience a relatively mild disease course with no symptomatic progression and limited involvement of major organs^4-7^. The factors that limit disease severity in ILE are unknown.

Genome-wide association studies have identified over 100 genes associated with SLE classification, including variants associated with specific disease manifestations, such as nephritis^8,9^. Increases in the number of these SLE-risk alleles, termed genetic load, are associated with SLE susceptibility^10-12^. Furthermore, increased genetic load correlates with more severe disease, organ damage, renal dysfunction, and mortality^13^. Therefore, we hypothesize that ILE may share genetic associations with SLE but with a reduced genetic load. However, the genetic risk of ILE has not been studied.

In this study, we determined the cumulative burden of SLE variants on ILE susceptibility by comparing the genetic load of SLE-risk alleles in European American (EA) ILE patients, SLE patients, and healthy controls.

## METHODS

### Study population

This study was performed in accordance with the Declaration of Helsinki and approved by the Oklahoma Medical Research Foundation (OMRF) Institutional Review Board. All participants provided written informed consent prior to study procedures. EA patients with SLE (n=171) or ILE (n=174) and healthy controls with no self-reported lupus manifestations (n=133) were selected from existing collections in the Arthritis & Clinical Immunology Biorepository (CAP # 9418302) at OMRF. Demographic information was self-reported. Participants with SLE or ILE were characterized by systematized medical records review for SLE classification criteria. ILE was defined as 3 ACR 1997 criteria and SLE as 4 or more ACR-1997 criteria^2^. Patients with ILE by ACR who also did not meet SLICC classification criteria^3^ were considered ILE^SLICC^. All individuals with ILE were previously enrolled in the Lupus Family Registry and Repository (LFRR) (1995–2012)^14^. Healthy controls with no documented lupus manifestations were also previously enrolled in the LFRR or from the Oklahoma Immune Cohort through the Oklahoma Rheumatic Disease Research Cores Center collections.

### Genotyping, quality control, and imputation

Samples were genotyped on the Infinium Global Screening Array (GSA)-24 v2.0 (Illumina, San Diego, CA), with 665,608 variants genotyped per sample. With consulting support from Rancho BioSciences (San Diego, CA), quality control was performed at the sample and variant level in PLINK 2.0 (v1.90) (Supplementary Figure 1). Samples with call rates below 90%, extreme heterozygosity measured by Wrights inbreeding coefficient (F< -0.05 || F > 0.1), or discordance between genotyped and clinically recorded sex were excluded. Variants from sex and mitochondrial chromosomes and somatic variants with minor allele frequency <0.1% were also excluded.

After quality control, 542,524 variants were available for imputation. The data were then pre-phased to infer underlying haplotypes with the 1000 Genomes phase III reference panel using SHAPEIT (v2.79), and whole-genome imputation was performed on the pre-phased haplotypes using IMPUTE (v2.3.2). To filter for variants of high imputation accuracy, only those with an information score >0.9 were retained.

### Genetic load

The genetic load was calculated for 478 subjects based on previously identified SLE-associated SNPs with genome-wide significance in the European population^11^. Of the 123 variants meeting tier 1 statistical significance (*P* > 5×10^−8^ and *P*FDR < 0.05)^11^, 99 met post-imputation quality control and were included for genetic load calculation (Supplementary Figure 1). Unweighted genetic loads were calculated as the total sum of risk alleles for each individual. Weighted genetic loads were defined as the sum of risk alleles multiplied by the beta coefficient (the natural logarithm of the previously published odds ratio [OR] of each risk allele for SLE susceptibility)^11^. If the beta coefficient was negative, the count for the reverse coded allele and the inverse OR was used.

### Statistical analysis

The genetic load was compared using Kruskal Wallis with Dunn’s post hoc test for multiple corrections. Statistical comparisons and receiver operator characteristic (ROC) analysis were performed using GraphPad Prism (v8.3.1). ORs were computed using Excel (v14.6.9), comparing individuals with a specific weighted genetic load (+/-2) to those within the lowest 10%. *P* values less than 0.05 were considered statistically significant.

## RESULTS

### ILE patients exhibit a similar increased SLE-risk allele genetic load as SLE patients

To assess the impact of known SLE genetic associations on ILE susceptibility, we compared the genetic load of a set of 99 previously described SLE risk variants^11^ in EA ILE patients (n=174), SLE patients (n=171), and unaffected controls (n=133) (Supplementary Table 1). Due to the low numbers of subjects from other races in the ILE cohort and challenges with combining race-specific genetic load information, we elected not to attempt any other race-specific genetic load comparisons. Consistent with previous findings^10-12^, EA SLE patients exhibited significantly greater unweighted and weighted genetic loads compared to healthy controls (Figure 1A and B). Unweighted and weighted genetic loads were also higher in EA ILE patients compared to healthy controls and did not differ compared to SLE patients (Figure 1A and B). We next stratified the ILE patients based on SLICC criteria, which is more sensitive compared to ACR criteria^3,15^. A similar trend was observed in ILE^SLICC^ patients (n=119) compared with SLE patients (Supplementary Figure 2A and B), suggesting a comparable genetic load in ILE and SLE patients irrespective of classification criteria used.

**Figure 1.**
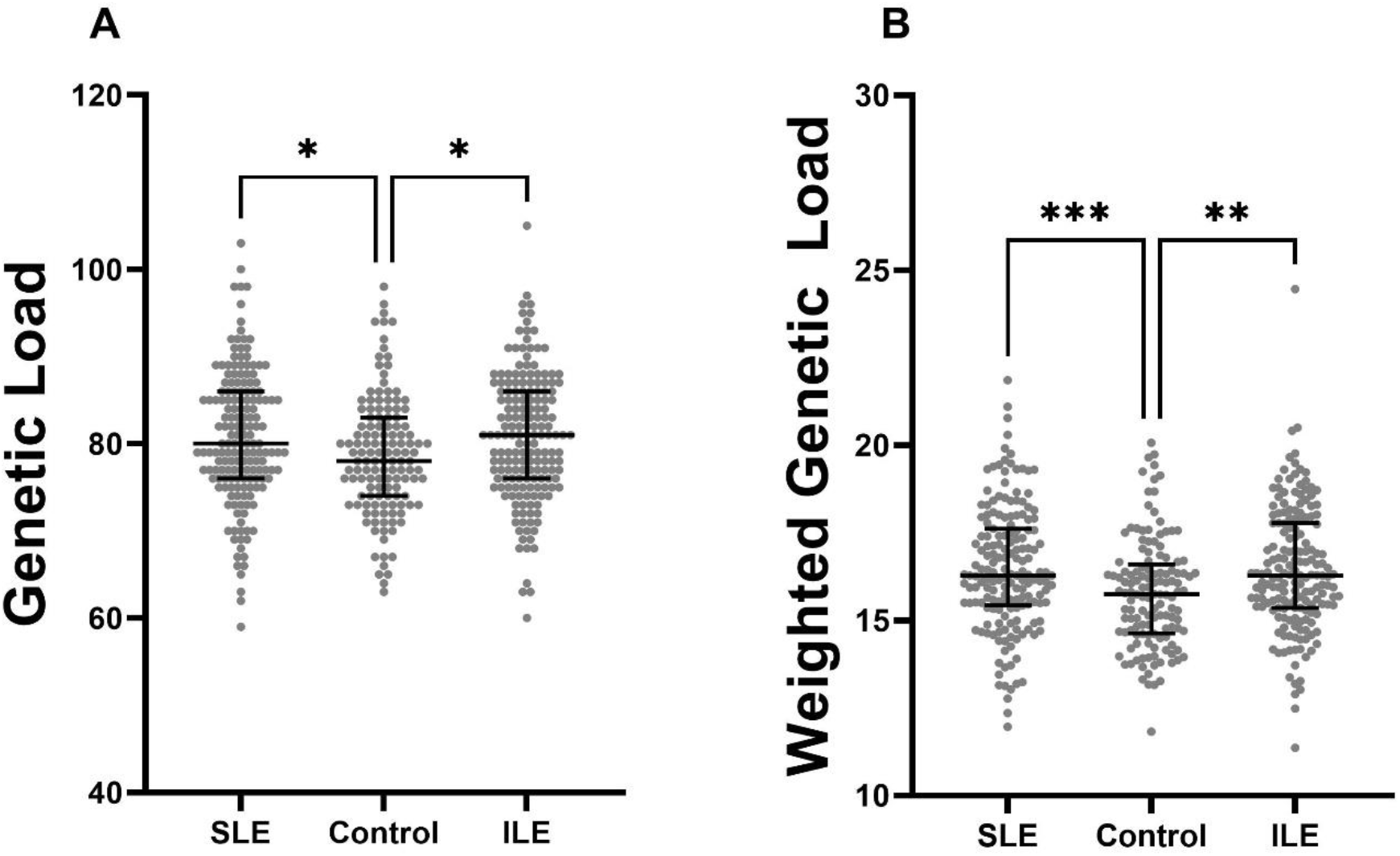
Incomplete lupus erythematosus (ILE) patients exhibit increased genetic load of systemic lupus erythematosus (SLE) risk alleles, similar to SLE patients. **(A)** Unweighted and **(B)** weighted SLE-risk allele genetic loads in European American SLE patients (n=171), ILE patients (n=174), and healthy controls (n=133). Graphs show the median and interquartile range. Statistical significance was determined using Kruskal Wallis with Dunn’s post hoc test. *p<0.05,**p<0.01, ***p< 0.001.

To understand how SLE-risk allele genetic load influenced the odds of disease in an individual, we calculated ORs comparing individuals with a given weighted genetic load (+/-2.0) to those within the lowest 10%. The probability of disease increased with increasing weighted genetic load for SLE, ILE, and ILE^SLICC^ patients compared to healthy controls (Figure 2A-C). Specifically, those with a weighted genetic load of 19 (+/-2.0) or higher showed greater odds of developing SLE or ILE compared to healthy controls (Figure 2A-C). However, the odds of developing SLE compared to ILE did not change with increasing weighted genetic load (Figure 2D). Similarly, higher genetic load differentiated ILE and SLE patients from controls (area under the curve = 0.62 for both), but not ILE patients from SLE patients (area under the curve = 0.51) by ROC analysis (Figure 2E). These results demonstrate that the milder phenotype of EA ILE is not due to a reduced dose of SLE genetic risk.

**Figure 2.**
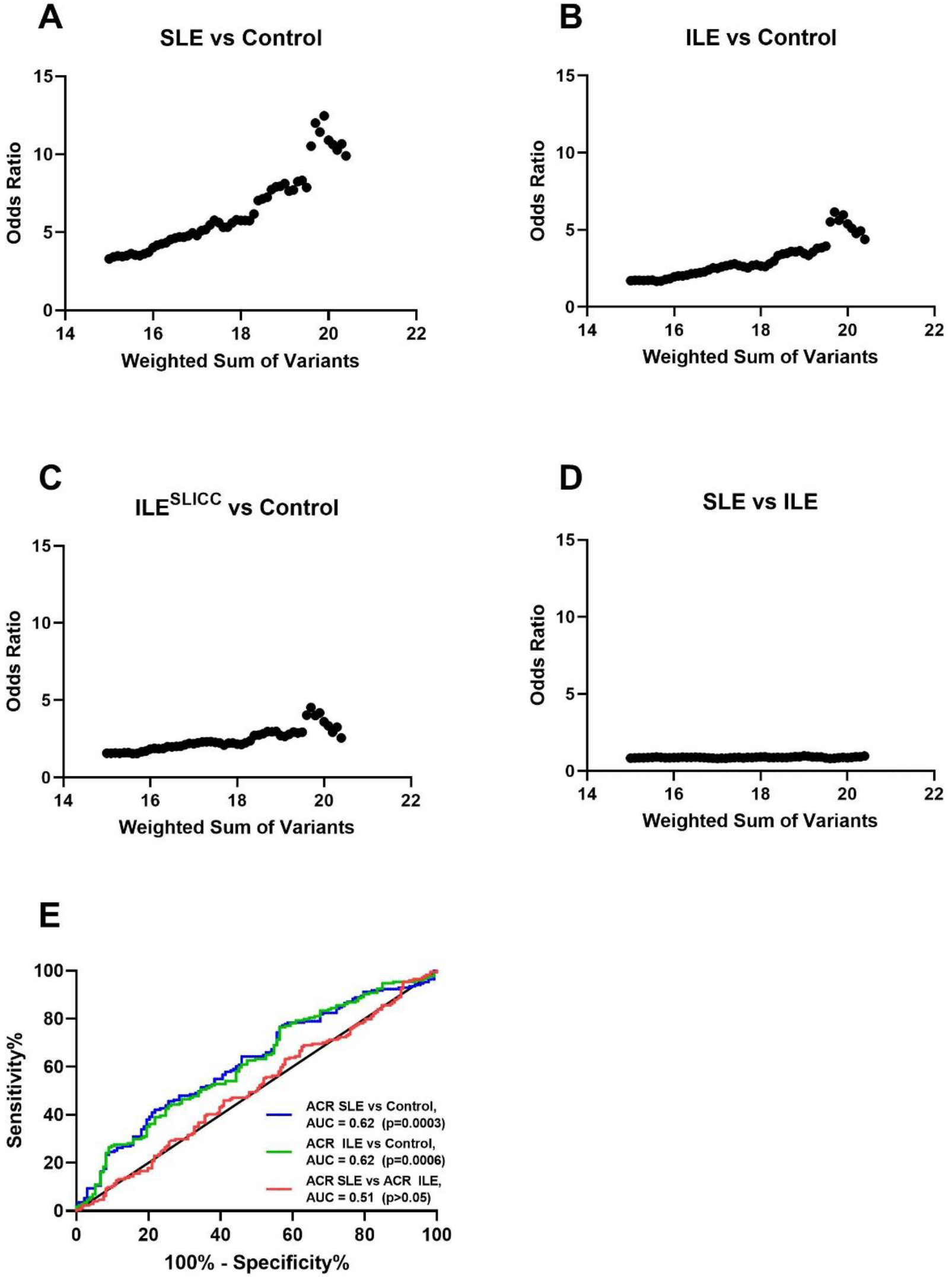
The genetic load of systemic lupus erythematosus (SLE) risk alleles does not distinguish incomplete lupus erythematosus (ILE) patients from SLE patients. **(A-D)** Odds ratios comparing individuals with a given weighted genetic load (+/-2.0) to those within the lowest 10% for **(A)** SLE patients (n=171) and healthy controls (n=133), **(B)** ILE patients (n=174) and healthy controls, **(C)** ILE patients who also do not meet SLICC criteria (ILE^SLICC^; n=119) and healthy controls, or **(D)** SLE and ILE patients. **(E)** Receiver operating characteristic analysis to assess the prediction ability of weighted genetic load in SLE patients, ILE patients, and healthy controls. AUC; area under the curve.

## DISCUSSION

This study is the first to determine the genetic load of SLE-risk alleles and unique risk variants in ILE. Although ILE patients exhibit a milder phenotype compared to SLE, the genetic load of SLE-risk alleles in ILE patients was indistinguishable from SLE patients, suggesting a similar genetic predisposition. However, it is unknown if there may be unique risk or protective variants associated with a subgroup of ILE patients who never progress to SLE classification.

Our study has some limitations. We were unable to examine race-specific genetic load differences between SLE, ILE, and healthy controls due to low numbers of subjects in the racial subgroups. Therefore, replication in larger race-matched cohorts and subsequent trans-ancestral meta-analysis is imperative.

Together, our data support an enhanced genetic predisposition towards ILE similar to SLE through aggregate genetic variants. Future studies in larger, longitudinal pre-clinical cohorts are needed to determine whether the phenotypic differences between SLE and ILE are governed by novel ILE genetic variants or disparate environmental or gene-environmental factors.

## ABBREVIATIONS

ACR: American College of Rheumatology
EA: European American
GSA: global screening array
ILE: incomplete lupus erythematosus
OMRF: Oklahoma Medical Research Foundation OR odds ratio
SLE: systemic lupus erythematosus
SLICC: Systemic Lupus International Collaborating Clinics

## CONFLICT OF INTEREST

There are no conflicts of interest to report.

## AUTHOR CONTRIBUTIONS

JMG, JAJ, MS conceived and designed the work. JAJ, TA captured, analyzed, and interpreted the patients’ data, while JMG, CM, MS, JK, SM, WD helped in experimental data acquisition. JMG, JAJ, MS, JK, MS contributed to data interpretation, and MS, JAJ, CAW, JMG contributed to the writing of the manuscript.

## FUNDING

These studies were supported by funds from the NIH which includes grants P30AR073750 (JAJ, JMG), U54GM104938 (JAJ), UM1AI144292 (JAJ, JMG), R01AR072401 (JAJ, JMG).

## ACKNOWLEDGMENTS

We would like to thank the Lupus Family Registry and Repository and OMRF Rheumatology Center of Excellence patients, controls, and staff. We also thank the OMRF Biorepository, OMRF Human Phenotyping Core, and OMRF Clinical Genomics Center for technical assistance. We would like to acknowledge the bioinformatics consulting support from Rancho BioSciences, especially their technical assistance in aspects relating to quality control of the genetic dataset, generation of the imputed, pre-phased data, and quality checks of these data that were subsequently used for calculating the genetic load values of the individuals in this study.

## Notes

### Competing Interest Statement

The authors have declared no competing interest.

