## Supplemental Information for "Genetic load in incomplete lupus erythematosus"

### Supplementary Material

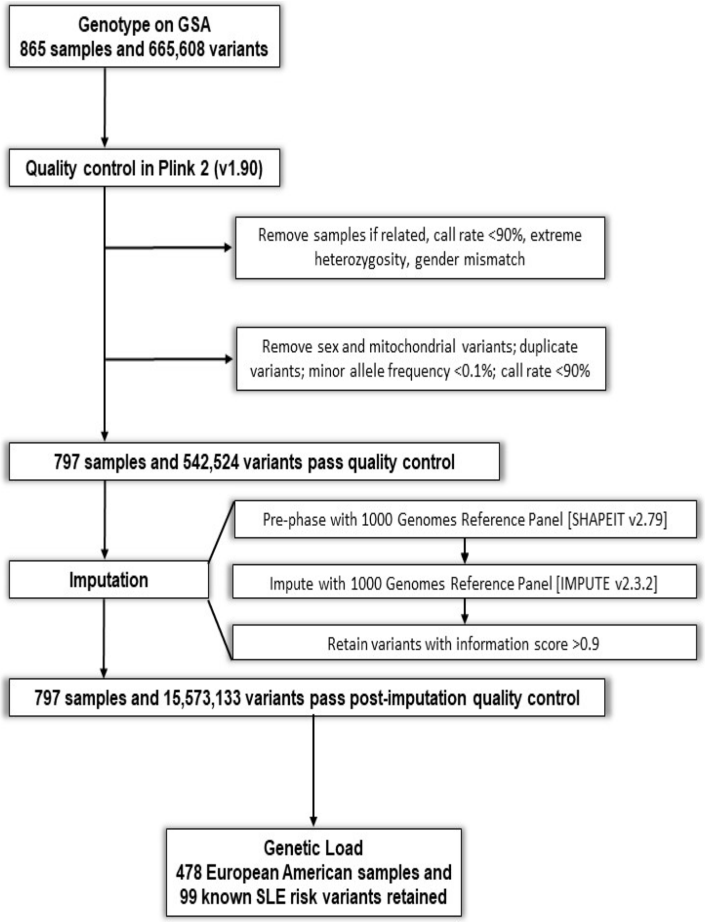


**Supplementary Figure 1. Genotyping and quality control pipeline**. 865 study participants were genotyped on the Illumina Global Screening Array (GSA) for genetic load calculations. Following a series of quality control steps, 478 European American (EA) patients were included for genetic load calculations.

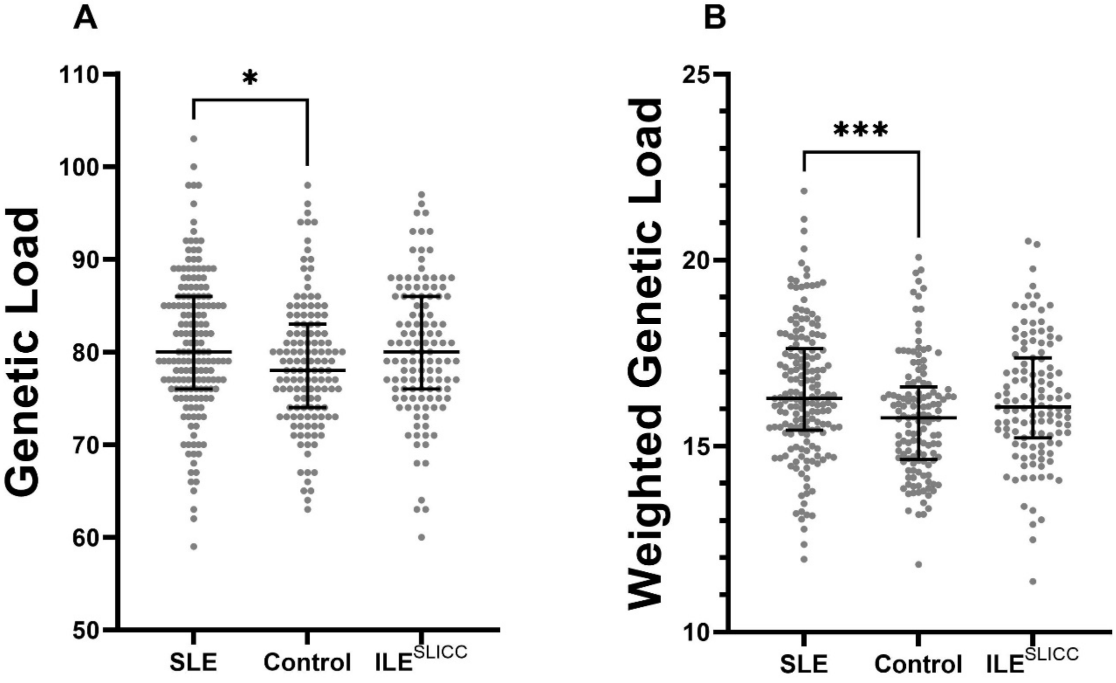


**Supplementary Figure 2. Incomplete lupus erythematosus (ILE) patients who do not meet SLICC criteria ($ILE^{SLICC}$) have a trend towards increased genetic load of systemic lupus erythematosus (SLE) risk alleles, comparable to SLE patients.** **(A)** Unweighted and **(B)** weighted SLE-risk allele genetic loads in European American SLE patients (n=171), $ILE^{SLICC}$ patients (n=119), and healthy controls (n=133). Graphs show the median and interquartile range. Statistical significance was determined using Kruskal Wallis with Dunn's posttest. *p<0.05, ***p< 0.001.

**Supplementary Table 1. Demographic characteristics of ILE and SLE patients and healthy controls used in genetic load calculations.**

| | **ILE**<br>(n=174) | **SLE**<br>(n=171) | **Control**<br>(n=133) |
|---|---|---|---|
| **Age,** mean (SD) | 47.3 (13.6) | 43.6 (13.7) | 46.6 (14.8) |
| **Female,** n (%) | 151 (87) | 152 (89) | 115 (86) |
